# Enhancers mapping uncovers phenotypic heterogeneity and evolution in patients with luminal breast cancer

**DOI:** 10.1101/193771

**Authors:** Darren K. Patten, Giacomo Corleone, Balázs Győrffy, Edina Erdős, Alina Saiakhova, Kate Goddard, Andrea Vingiani, Sami Shousha, Lőrinc Sándor Pongor, Dimitri J. Hadjiminas, Gaia Schiavon, Peter Barry, Carlo Palmieri, Raul C. Coombes, Peter Scacheri, Giancarlo Pruneri, Luca Magnani

## Abstract

The degree of intrinsic and interpatient phenotypic heterogeneity and its role in tumour evolution is poorly understood. Phenotypic divergence can be achieved via the inheritance of alternative transcriptional programs^1,2^. Cell-type specific transcription is maintained through the activation of epigenetically-defined regulatory regions including promoters and enhancers^1,3,4^. In this work, we annotated the epigenome of 47 primary and metastatic oestrogen-receptor (ERα)-positive breast cancer specimens from clinical samples, and developed strategies to deduce phenotypic heterogeneity from the regulatory landscape, identifying key regulatory elements commonly shared across patients. Highly shared regions contain a unique set of regulatory information including the motif for the transcription factor YY1. *In vitro* work shows that YY1 is essential for ERα transcriptional activity and defines the critical subset of functional ERα binding sites driving tumor growth in most luminal patients. YY1 also control the expression of genes that mediate resistance to endocrine treatment. Finally, we show that H3K27ac levels at active enhancer elements can be used as a surrogate of intra-tumor phenotypic heterogeneity, and to track expansion and contraction of phenotypic subpopulations throughout breast cancer progression. Tracking YY1 and SLC9A3R1 positive clones in primary and metastatic lesions, we show that endocrine therapies drive the expansion of phenotypic clones originally underrepresented at diagnosis. Collectively, our data show that epigenetic mechanisms significantly contribute to phenotypic heterogeneity and evolution in systemically treated breast cancer patients.

## Introduction

Breast cancer (BC) is the most common cancer type and the second most frequent cause of cancer related death in women^5^. 70% of all BC cases contain variable amounts of oestrogen receptor-alpha (ERα) positive cells. ERα is central to BC pathogenesis and serves as the target of endocrine therapies (ET)^6,7^. ERα-positive BC is typically subdivided in two ‘intrinsic’ molecular subtypes (luminal A and luminal B^8^) characterized by distinct prognosis, highlighting functional inter-patient heterogeneity. Recent analyses demonstrate that patient-to-patient heterogeneity is more pervasive (reflected by histological^9^, genetic architecture^10^ and transcriptional^11^ differences) ultimately influencing long-term response to endocrine treatment^12^. Indeed, 30-40% of ERα BC patients relapse during or after completion of adjuvant endocrine therapies. At the time of relapse, almost all patients will have developed resistance to ET, partly through treatment-specific genetic evolutionary trajectories^13^. Additionally, the presence of genetic intra-tumor heterogeneity has also now been extensively documented in several cancer types, demonstrating the role of clonal evolution in cancer ^14^. Yet, recent studies have shown that driver coding-mutations do not significantly change between primary and metastatic luminal breast cancer, with the notable exception of *ESR1* mutations^15^, suggesting that alternative mechanisms might contribute to BC progression and drug-resistance. Parallel to genetic evolution, phenotypic/functional changes driven by epigenetic mechanisms can also contribute to breast cancer progression and ET resistance in cell lines^16,17^. Nevertheless, little is known about the epigenome of BC patients, its influence on intra-tumour phenotypic heterogeneity and its role in breast cancer progression.

Epigenetic modifications consist of chemical modifications targeting the DNA/RNA (e.g. DNA methylation) and the chromatin (histone modifications). Histone modifications have been successfully used to map regulatory regions and to annotate the non-coding DNA^1,3^. Acetylation of lysine 27 on histone 3 (H3K27ac) is strongly associated with promoters and enhancers of transcriptionally active genes^4,18,19^. Increasing evidence suggests that epigenetic information can actively transfer gene transcription states across cell division^20-23^. Epigenetic modifications play also a central role in modulating ERα binding to the DNA possibly by interacting with ERα-associated pioneer factors ^24,25^. Finally, *in vitro* studies have shown that epigenetic reprogramming might play a central role in ERα BC cells that adapt to endocrine therapy^16,17^.

Here we show the results of a systematic investigation of the epigenetic landscape of ERα positive primary and metastatic breast cancer from 47 individuals. Our results represent the first large scale topographic mapping of the active regulatory landscape of longitudinal ERα-positive BC. Using H3K27ac we mapped active promoters and enhancers across treatment naïve primary and endocrine treated metastatic patients. We used bioinformatic approaches to deconvolute the complex regulatory landscape and identified inter- and intra-patient epigenetic heterogeneity. We mined promoters and enhancers from clinically relevant breast cancer samples for potential regulatory drivers identifying YY1 as a novel key player in ERα-positive BC. Finally, we demonstrate that epigenetic mapping can efficiently estimate phenotypic heterogeneity changes throughout BC progression.

## Results

### Mapping enhancers and promoters of primary and metastatic ERα positive breast cancer

To build a comprehensive compendium of all the clinically relevant active regulatory regions of luminal BC we profiled fifty-five ERα positive BC samples (primary n=39, and metastatic n=16) with H3K27ac ChIP-seq (Supplementary Table S1). To minimize the introduction of noise from non-tumor tissues we used samples with high tumor burden (>70%, Supplementary Figures S1). 85% of samples yielded satisfactory results (47/55, Supplementary Figure 2A and Table S2). H3K27ac-enriched regions were classified into 23,976 gene-proximal (1kb upstream of transcription start site (TSS), promoters) and 326,719 gene-distal (enhancers). Considering the ten-fold difference in H3K27ac signal, it was not surprising to observe that 80% of promoters can be captured by profiling 4 individual, while nearly 40 are needed to reach the same coverage for enhancers, as indicated by saturation plots (Supplementary Figure 2B). These data are in agreement with enhancers being the main determinants of cell-type specific transcriptional differences ^4,18,26,27^. To gain insights on the penetrance of each regulatory region, we developed a Sharing Index (SI) by annotating all enhancers and promoters in function of the number of patients sharing the H3K27ac signal at each specific location (Supplementary Figure 2C). In agreement with saturation analyses, we find that a large portion of enhancers are patient-specific (SI=1) while active promoters are more commonly shared between patients (Supplementary Figure 2C). Collectively, these data demonstrate that enhancers account for the majority of potential epigenetic heterogeneity in ERα-positive BC.

### Enhancer activity allows the qualitative assessment of phenotypic heterogeneity

Genetic heterogeneity is an hallmark of most solid tumours ^28^. Nonetheless, genetic intra-tumoral heterogeneity does not often directly translates to phenotypic heterogeneity. In agreement, despite extensive inter- and intratumoral clonal genetic diversity^29^, the majority of ERα-positive tumors benefit from systemic ET^12^. Likewise, treatment-naïve metastatic patients generally respond well to ET, at least initially, suggesting that genetic heterogeneity on its own cannot explain treatment resistance/response. On the other hand, phenotypic hierarchies can override genetic hierarchies in brain cancers ^2,30^, suggesting that inheritable epigenetic program might ultimately contribute to phenotypic heterogeneity and treatment outcome.

The existence of intra-tumoral phenotypic heterogeneity in breast cancer patients has been known to pathologists for decades, at least for a small number of biomarkers. For example, immunohistochemistry (IHC) assessment of the proportion of ERα-positive within luminal cancer patients varies on a *continuum* from less than 1% to nearly 100%^31^. Unfortunately, IHC is low-throughput method, and typically only a few proteins can be studied before the sample is consumed. In contrast, assessing phenotypical heterogeneity from patient bulk transcriptional data is unfeasible as transcription is ultimately an analogue signal in which each individual cell can contribute a stochastic amount of RNA, making data deconvolution impractical (Fig 1A). For instance, cells with focal gene amplification have higher bulk gene expression but individual cells can contribute radically different amounts as shown by single-molecule single-cell RNA FISH^13^. On the other hand, recent evidence show that the signal captured by chromatin assays such as ATAC-seq appears to be directly proportional to the cells contributing to it^32^. Similarly, ChIP-seq signal can be thought of as digital information with each single nucleosome being ON (K27ac) or OFF at any given time (Fig. 1A). Notably, even within genetically clonal cell lines, the H3K27ac signal varies considerably between different regulatory regions. Regulatory regions labelled as super enhancers, for example, have 10-100-times more H3K27ac signal than typical enhancers^18^. What accounts for the variation in signal is not known, but one possibility is that heterogeneity within the cell population (either clonal or sub-clonal) contribute to signal intensity. For example, super-enhancers might represent regulatory regions active across most or all cells within a population at any given time (clonal, H-peaks), while “typical” enhancers with lower H3K27ac signal may represent sub-clones (M and L peaks, Fig. 1A). This concept is similar to measuring variant allele frequencies (VAF) to infer genetic heterogeneity. Phenotypical heterogeneity might be the consequence of heterogeneous cell populations (i.e. tumor, stroma and immune infiltrate) or actual cancer-specific epigenetic subclones. As our ChIP-seq data are derived from high tumor burden samples, we hypothesized that H3K27ac signal could allow for a qualitative assessment of phenotypic heterogeneity. We further theorised that direct correlation might exist between clonal prevalence (intra-tumour) and population prevalence (inter-patients) (Fig. 1A).

**Figure 1:**
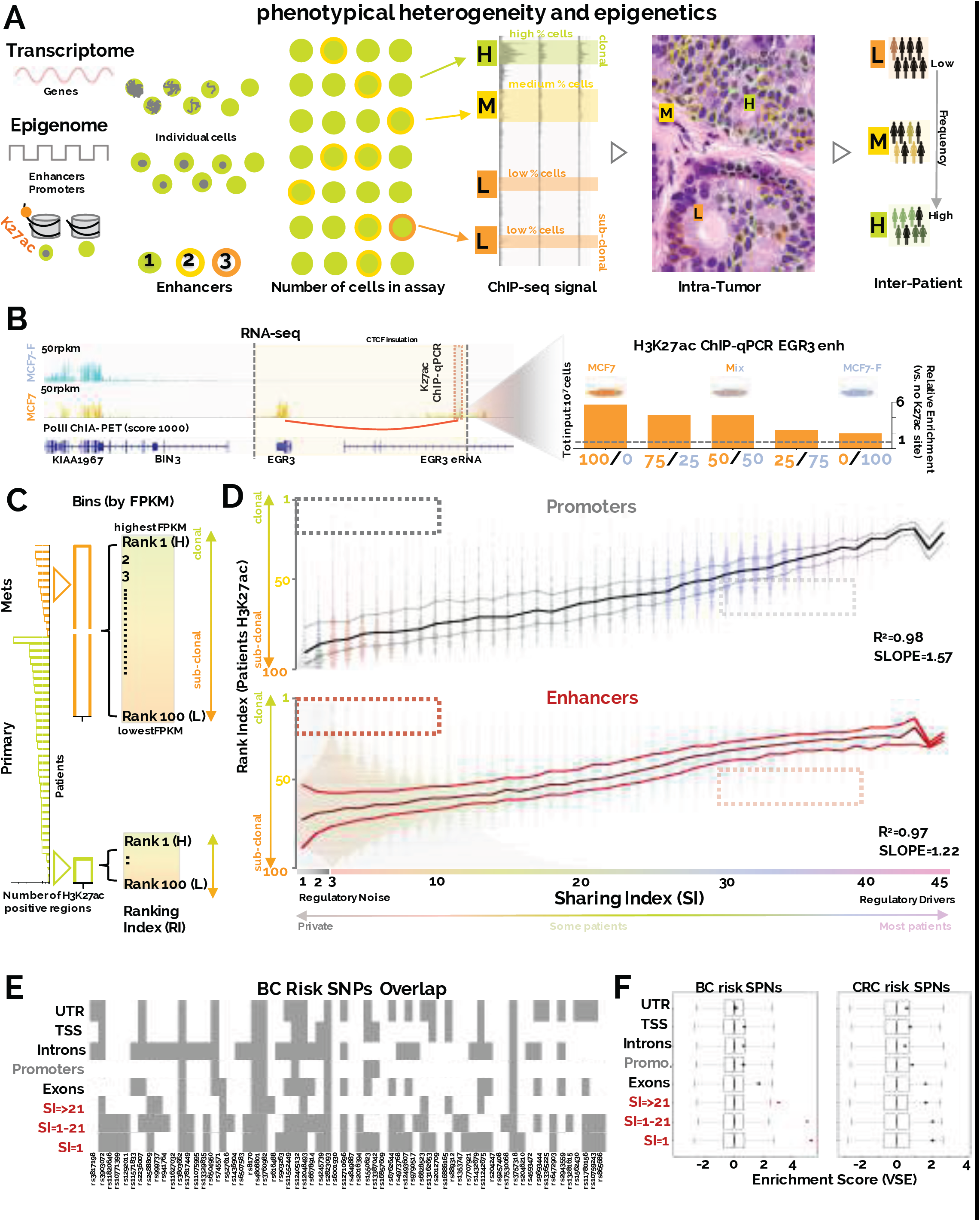
Assessment of inter-and intra-tumor epigenetic heterogeneity. A) Main hypothesis of the study. Transcriptional data from bulk tissue represent the average over million cells. Each cell contributes a value from a continuous distribution of potential mRNA molecules. For chromatin data, each cell can only contribute a deterministic value to the bulk signal, generally from two alleles. Therefore, the relative strength of ChIP-seq data is dependent on the number of cells carrying epigenetic signal at discrete loci. H, M and L represent strong, medium and weak signal, respectively. Clonal regulatory regions are commonly shared by BC patients while weak enhancers are more patient specific B) EGR3 mRNA is expressed in MCF7 but not MCF7-F cells. eRNA and Pol-II ChIA-PET show enhancer activity in MC7 but not MCF7-F^16^. CTCF insulated perimeter is shown in yellow. Predicted looping from ChIA-PTE is shown in red. The observed ChIP-qPCR signal for H3K27ac at EGR3 enhancers decrease with increasing number of MCF7-F cells mixed in the sample C) Ranking strategy: H3K27ac signal is normalized at each locus and assigned a ranking index based on relative strength within each single ChIP-seq experiment (1=strongest, 100=weakest, binning on RPKM signal). Binning is repeated for each patient. D) Linear regression shows that clonal enhancers are commonly shared between breast cancer patients. Y axis=Ranking Index, X axis=Sharing index. Sharing index indicate the number of patients sharing the regulatory region. Each dot represents the median RI (all patients) for a single regulatory region. The interpolating lines represent the median RI value and interquartile ranges for regulatory regions with the same SI E) Overlap between BC risk variants and annotated DNA elements F) Variant Set Enrichment analysis indicates that BC-specific but not CRC-specific GWAS risk variants occur more frequently than expected within the enhancers elements identified in our study.

We tested the initial assumption by performing spike-in experiments in which known numbers of cells with well-characterized regulatory region activity (and similar genetic background)^16^ were admixed in incremental proportions prior to H3K27ac ChIP-qPCR. The data shows that H3K27ac enrichment is proportional to the number of cells with the active enhancer (Fig. 1B). These findings are corroborated by an independent analysis using a different antibody (ERα) (Supplementary Figure S3A). As the signal between different patients is not directly comparable, we normalized the data using a ranking approach, assigning to each H3K27ac signal a Rank Index (RI, 1 to 100, strongest to weakest) (Fig. 1C). Signal from low RI (H peaks) might be associated with clonal regulatory regions active in almost all cells. Conversely, high RI (M-L peaks) mark more heterogeneous/sub-clonal enhancer activity. By investigating the relationship between RI and SI we found extremely high correlation between these two parameters (Fig. 1D), suggesting that clonal regulatory regions are more common between patients while sub-clonal regulatory elements are more patient-specific. We defined clonal low-RI/high-SI loci as regulatory drivers (RD, SI>21) and high-RI/low-SI that might originate from sub-clonal populations as regulatory noise (RN, SI<21). Of note, a small but discrete proportion of promoters/enhancers escape this general trend having extremely low RI despite being patient-specific or higher RI while being shared (dot-boxes, Fig. 1D).

### Enhancers are associated with BC risk-SNP and control gene transcription

We next investigated the extent to which regulatory regions identified in our cohort associate to BC. Previous analyses from ERα-BC cell lines, have shown that genetic predisposition might occur through SNPs that modulate transcription factors binding at enhancers (FOXA1 and ERα^33^). We then tested the relationship between DNA risk variants specifically associated with BC through GWAS^33,34^ and regulatory regions captured in patients. Strikingly, almost the totality of validated BC risk variants is contained within our H3K27ac database. Currently, this dataset represents the most enriched annotation for GWAS variants in breast cancer (Fig 1E). This overlap is highly significant specifically for enhancers but not for other annotations (Fig. 1F). Notably, this association is not replicated using colorectal cancer risk variants suggesting that these enhancers might play a specific role in BC development (Fig. 1F).

Next, we assessed the relationship between estimated enhancers clonality and transcriptional output. Transcriptional data obtained using microarray and RNA-seq estimate the average expression level within a population. The average expression is function of the number of cells engaged in active transcription and the number of RNA molecule within each cell^35^. Interestingly, several lines of evidence suggest that RNA transcription is stochastic ^13,36^, thus implying that the total number of cells with active transcription significantly contribute to changes in average RNA levels in bulk populations. As our analysis allows for qualitatively prediction of clonality in enhancer activity, it allows to test if clonal enhancers active in the majority of cells correlate with higher RNA levels. To do so we linked enhancers to their potential target genes using CTCF insulated boundaries^37^. We then analysed three independent BC expression datasets (METABRIC ^10^, TCGA ^38^ and Affymetrix ^39^) in function of RI/SI indexes. Our analyses show a predictable increase for mRNA levels with parallel increases in the associated SI, further suggesting that RDs drive RNA expression in a progressively increasing number of cells (Supplementary Figure 3B). These results were more modest when analysing the transcriptome from normal breast tissue (Supplementary Figure 3B, small insets) suggesting that our analysis has identified a subset of regulatory regions strongly associated with malignant outgrowth. These data indicate that transcripts identified as dis-regulated in BC might reflect changes in the size of phenotypic subpopulations. This could be in part driven by a selection process, as normal breast transcriptional data reflect the heterogeneous composition of the tissue (adipocyte, myoepithelial and epithelial cells), while BC is normally dominated by epithelial morphology. Collectively, our data show that enhancer activity strongly tracks transcriptional changes in breast cancer patients.

### Imputed transcription factors landscape of ERα breast cancer patients

Enhancers stores regulatory information in the form of transcription factors (TFs) binding motifs^40^. The vast majority of TFs require accessible chromatin in order to bind their cognate DNA sequences ^41^. We reasoned that a systematic investigation of the predicted TFs landscape in function of enhancer activity could reveal potential transcriptional BC drivers. To narrow down to the accessible DNA within active enhancers and promoters we integrated the DNaseI signal from 129 cell lines with the inferred nucleosome pattern obtained from the H3K27ac data (Fig. 2A). Initial analyses collapsing all imputed DHS in relationship with their enhancers and promoter location identified correctly well-known BC-TFs according to their promoter–enhancer bias (Supplementary Figure 4). We then stratified the complete set of enhancers and promoter regions based on the associated SI and repeated TF motif analysis focusing within each SI-defined bin followed by unsupervised clustering. This analysis generated two major clades (Fig. 2B), indicating the presence of different classes of regulatory regions. Strikingly, we find that RD and RN enhancers and promoters cluster specifically into the two major clades, suggesting that putative clonal and sub-clonal enhancers contain distinct regulatory information (Fig. 2B). Functional TF binding is associated with TF leaving a footprint within chromatin accessible regions ^40,42^. Interestingly, clonal enhancers in ERα-positive MCF7 breast cancer cells are significantly enriched in TF footprints^16^, while sub-clonal enhancers are significantly deprived of footprints suggesting that TFs might bind clonal enhancers with longer residence time ^42^ (Fig. 2C).

**Figure 2:**
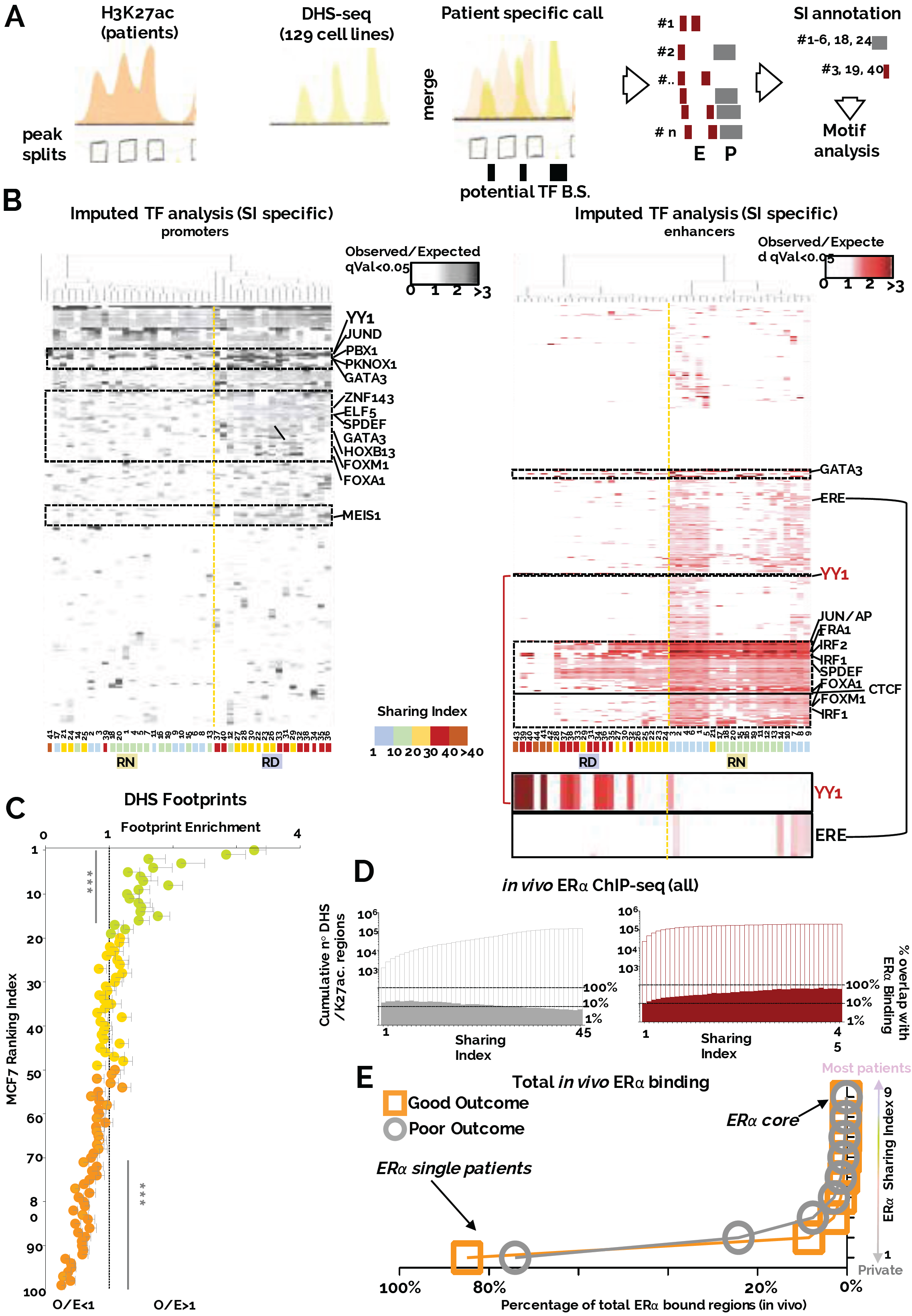
Clonal and sub-clonal regulatory regions contain distinct regulatory information. A) Bioinformatic framework of the analyses. H3K27ac calls were split to identify approximate nucleosome-level enrichment (sub-peaks). Sub-peaks data were integrated with ENCODE-derived DHS-seq calls to identify potential sites of TF binding. Individual imputed DHS regions were assigned SI values based on the number of patient sharing the region B) Transcription factor motif analysis of individual bins (SI) followed by unsupervised clustering. RD and RN regions cluster separately in two distinct clades. ERE and YY1 motif are blown up at the bottom C) Clonal enhancers in MCF7 cells (RI<20) are characterized by a higher number of TF footprints, while sub-clonal enhancers (RI>70) have less footprint than expected (O/E=1). Asterisks represent a pValue of <0.001 in a Wilcoxon Signed Rank Test D) Overlap of imputed DHS regions with *in vivo* derived ERα binding sites. The left Y axis indicates cumulative DHS regions. The right Y axes indicate the percentage of overlap based on total DHS in each SI bin E) Distribution plot of *in vivo* derived ER binding sites versus the number of patients in which they were observed^43^.

We then focused on estrogen-response elements (ERE) as they constitute the canonical DNA sequence to which ERα binds. Unexpectedly, ERE motifs are enriched only in RN enhancers, suggesting that a significant amount of ERα binding occurs in sub-clonal/less functional enhancers (Fig. 2B). To gain further insights on ERα dynamics we turned to a recently published ERα dataset obtained from patient material (n=15) ^43^. Generally, the proportion of ERα binding sites overlapping enhancers increase with the SI (9% vs. 70%, SI-1 vs. SI-39, Fig. 2C). This was not observed for promoters (15% vs. 6%, SI-1 vs. SI-39, Fig. 2C), and is consistent with previous studies demonstrating a significant bias for ERα binding at active enhancer elements^44,45^. These data imply that shared enhancers have a strong propensity for ERα binding despite being generally under-enriched in EREs (Fig. 2B). More importantly, the bulk of ERα binding were captured only once in fifteen patients (ERα SI=1), with less than 0.003% of ERα being shared across 75% of the patients (484 core ERα)^43^ (Fig. 2D). Together, these data support biochemical evidence that suggest that only a small fraction of ERα binding events with longer-residency time is functional^42^. We therefore concluded that the largest portion of ERα binding identified in patients occur at patient specific, sub-clonal enhancers and might be the consequence of the transient ERα-DNA interactions occurring while the receptor scans the genome^42^. The discrepancy between the small number functional ERα core binding and the observation of ERE-poor RD enhancers led us to hypothesize that other TFs might collaborate with ERα to increase its transcriptional efficiency at clonal enhancer. Most TF motifs enriched in RD enhancers are also largely observed in RN regions, with the notable exception of the motif for the transcription factor YY1 (Fig. 2B). Interestingly, TF analysis of the footprints within MCF7 clonal enhancer (Fig. 2C, RI<20) similarly identifies YY1 as the top hit (qVal=0.001). YY1 has been recently implied in *de novo* formation of enhancer promoter looping during neural development ^46^ and MYC-like ability to potentiate gene expression^47^ indicating a potential role in modulating the enhancer landscape in ERα-positive BC.

### YY1 enhancer activity mark a dominant phenotypic clone in BC

YY1 is a ubiquitously expressed TF (Supplementary Figure 5A-B) that can act as an activator or repressor by binding DNA, RNA and chromatin modifiers^48,49^. YY1 function in breast cancer is poorly understood, as it has been linked to different outcomes depending on breast cancer cells subtypes. In luminal BC, YY1 appears to be positively correlated with AP2 to promote HER2 activity ^50^, while in triple negative BC models appears to be a tumor suppressor by controlling BRCA1 expression ^51^. Interestingly, YY1 drosophila homolog PhoRC is involved in epigenetic memory by recruiting of Polycomb repressor complex to sequence specific regions^52^, but YY1’s role in mammals is not entirely understood ^46,47,53^. Our TF analysis shows that YY1 might actually operate as a global reader of active clonal enhancers common to the majority of ERα-positive patients. These data therefore predict that most luminal breast cancers should contain a dominant YY1-positive clone. To assess the size of YY1 phenotypic clone in our patient’s dataset we identified the *bona fide* enhancers looping at YY1 promoter using 3D chromatin maps ^54^ (Supplementary Fig. 6A). We found three potential enhancers with high SI within a CTCF-insulated region with YY1 promoter (SI A=41, B=33 and C=26, Supplementary Fig. 6A). Interestingly Enhancer A also directly interacts with Enhancer B-C, suggesting a multi-enhancers interaction with YY1 promoter. Enhancer A consistently ranks among the most clonal enhancer in nearly all patients, suggesting that YY1 is transcribed in almost all cells (Fig. 3A). By comparison, in most normal tissues profiled by H3K27ac within the Epigenome Roadmap consortium^45^, YY1 Enhancer A activity is more variable while remaining relatively dominant, implying that some tissues may harbour YY1-subclonal subpopulations (Fig 3B). Consistent with these predictions, immunocytochemistry (IHC) meta-analysis (Fig 3B) showed a decreasing number of YY1 positive cells in correspondence to increasing RI scores (Insets, Fig 3B and Supplementary Figure 6B). Collectively, these data suggest that enhancer ranking can capture qualitative changes in intra-tumoral heterogeneity, and that YY1-enhancer activity marks a dominant phenotypic clone in ERα-positive BC.

**Figure 3:**
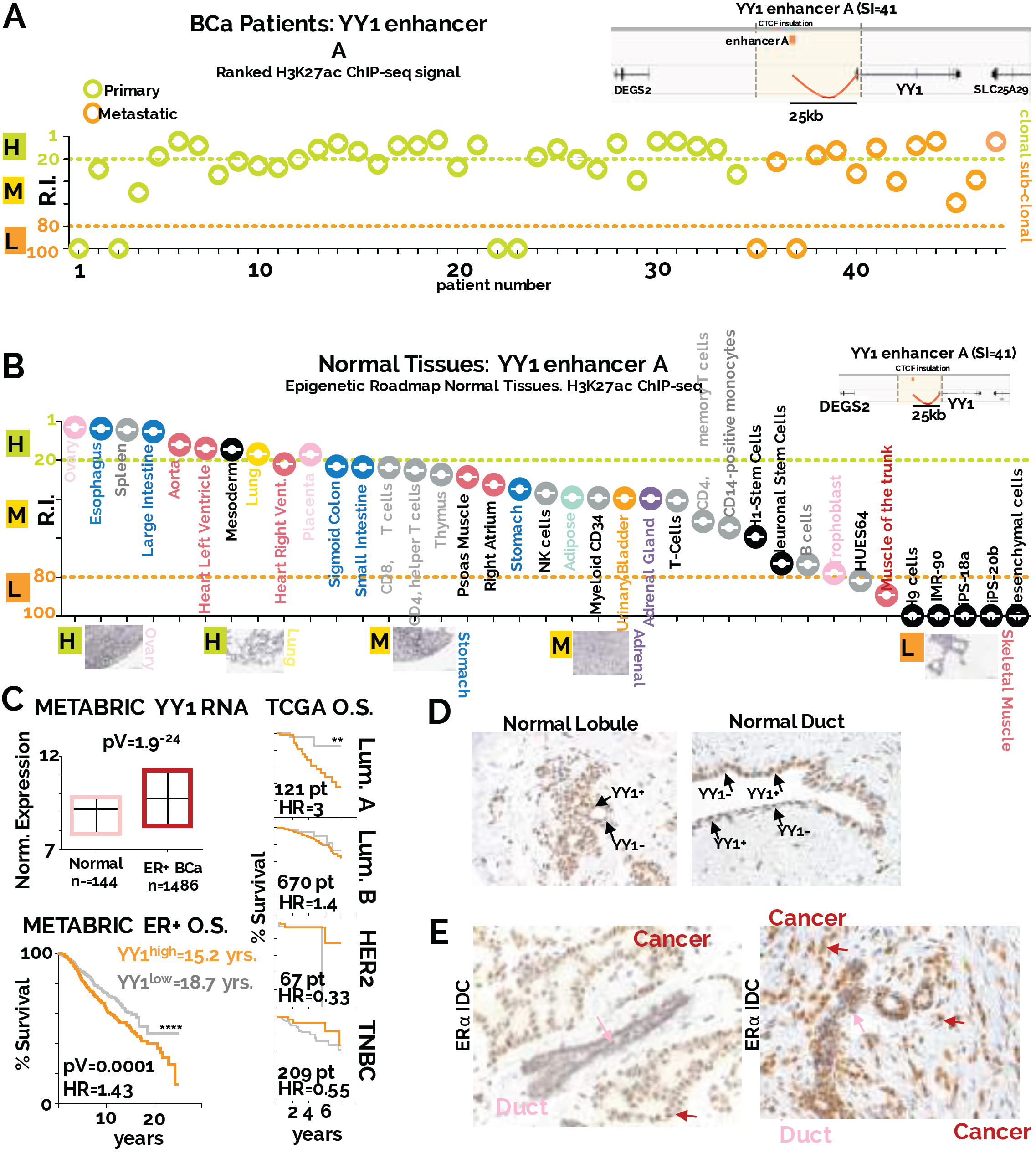
YY1 identify a dominant phenotypic clone in ER BC. A) RIs for the YY1 enhancer within all the individual patients included in the current study. YY1 enhancer location with its 3D interactions are shown in the top right inset B) YY1 enhancer ranking analysis of available Epigenome Roadmap H3K27ac datasets. Tissues are displayed from the strongest to the weakest YY1 enhancer activity (based on RI). Representative IHC analysis of normal tissues stained with a YY1 antibody are shown C) Top left: YY1 expression in ER-positive breast cancer compared to normal breast tissue. Bottom left: Kaplan-Meier analysis of patient outcome using YY1 expression to stratify patients. Right: Kaplan-Meier analysis of patient outcome using YY1 expression. All BC subtypes were analysed separately D) IHC analysis of normal breast tissues highlights YY1 functional subclones in normal breast E) IHC analysis of ER positive invasive ductal carcinomas identify YY1 positive clones as the dominant clonal population.

Next, we looked at the significance of YY1 mRNA expression in a pan-cancer analysis and found that tumor tissue generally have significantly higher expression level for YY1 as compared to normal tissues (Supplementary Figure 7A). This does not appear to reflect simply the proliferation status as we found no correlation between YY1 and Ki67 expression in 2509 Breast Cancer patients ^55^. This observation was replicated in an independent large BC dataset as well (Fig. 3C and Supplementary Figure 7B). Of note, YY1 is not subject to recurrent genomic aberrations (data not show). These data suggest that cancer lesions might contain a larger fraction of YY1-expressing cells as compared to more heterogeneous tissues (Fig. 3B). Meta-analysis of the METABRIC^10^ datasets shows that patients with higher YY1 mRNA level at diagnosis have significantly worse outcome (Fig. 3C). We replicated this finding using TCGA RNA-seq data with significant stratification occurring in Luminal A patients, a subtype typically associated with good prognosis (Fig. 3C). To test if YY1 increased mRNA expression could be driven by an expansion of YY1-positive cells from a more heterogeneous population, we stained normal breast section with IHC. Our data show that lobules and ducts contain distinct YY1 positive sub-clonal populations within the luminal and basal compartments (Fig 3D). On the other hand, nearby tumour tissue is overwhelmingly YY1 positive, demonstrating the existence of a clonal YY1 population. The data demonstrate that the expansion of a YY1 phenotypic clone drive the changes in bulk RNA levels between normal and tumor samples and reinforce the notion that YY1 might play a central role in ERα-positive BC.

### YY1 is a global enhancers modulator marking functional ERα binding

The TF analysis in our epigenomic patient dataset revealed YY1 uniquely associated with clonal regulatory regions and potentially collaborating with ERα at critical enhancers. To gain mechanistic insight on the role of YY1 we performed YY1 ChIP-Seq in quiescent (estrogen-deprived) and estrogen-stimulated luminal BC MCF7 cells. Cells were stimulated with estrogen for 45 minutes, upon which maximum ERα-binding to chromatin occurs^45^. ChIP-seq biological replicates show very high correlation (R^2^=0.98), thus we kept consensus loci for further analyses. In quiescent cells, YY1 occupies a very small set of enhancers and promoters near housekeeping genes^56^ (Fig. 4A). Strikingly, estrogen stimulation induce a 23-fold expansion of the YY1 binding repertoire, mostly at enhancer regions (Fig. 4A). Newly occupied loci are associated with ERα-BC signatures and epigenetic editors (Fig. 4A). Interestingly, only ∼10% of all binding is characterized by a high affinity YY1 motif suggesting that induced YY1 could also bind directly to modified nucleosomes through its chromatin remodelling partner INO80^57^. Orthogonal analyses show that induced-YY1 binding involves almost all MCF7 active regulatory regions and is strongly associated with H3K27ac marks (Fig. 4B). Nonetheless, it is unlikely that YY1 binding directly promote or requires H3K27ac as we did not find any difference between quiescent (estrogen-deprived) and estrogen-stimulated H3K27ac epigenomes (data not shown). Conversely, YY1 binding is absent from silenced genes (Supplementary Figure 7C), demonstrating that YY1 does not associate with PRC2 mediated repression in BC cancer cells.

**Figure 4:**
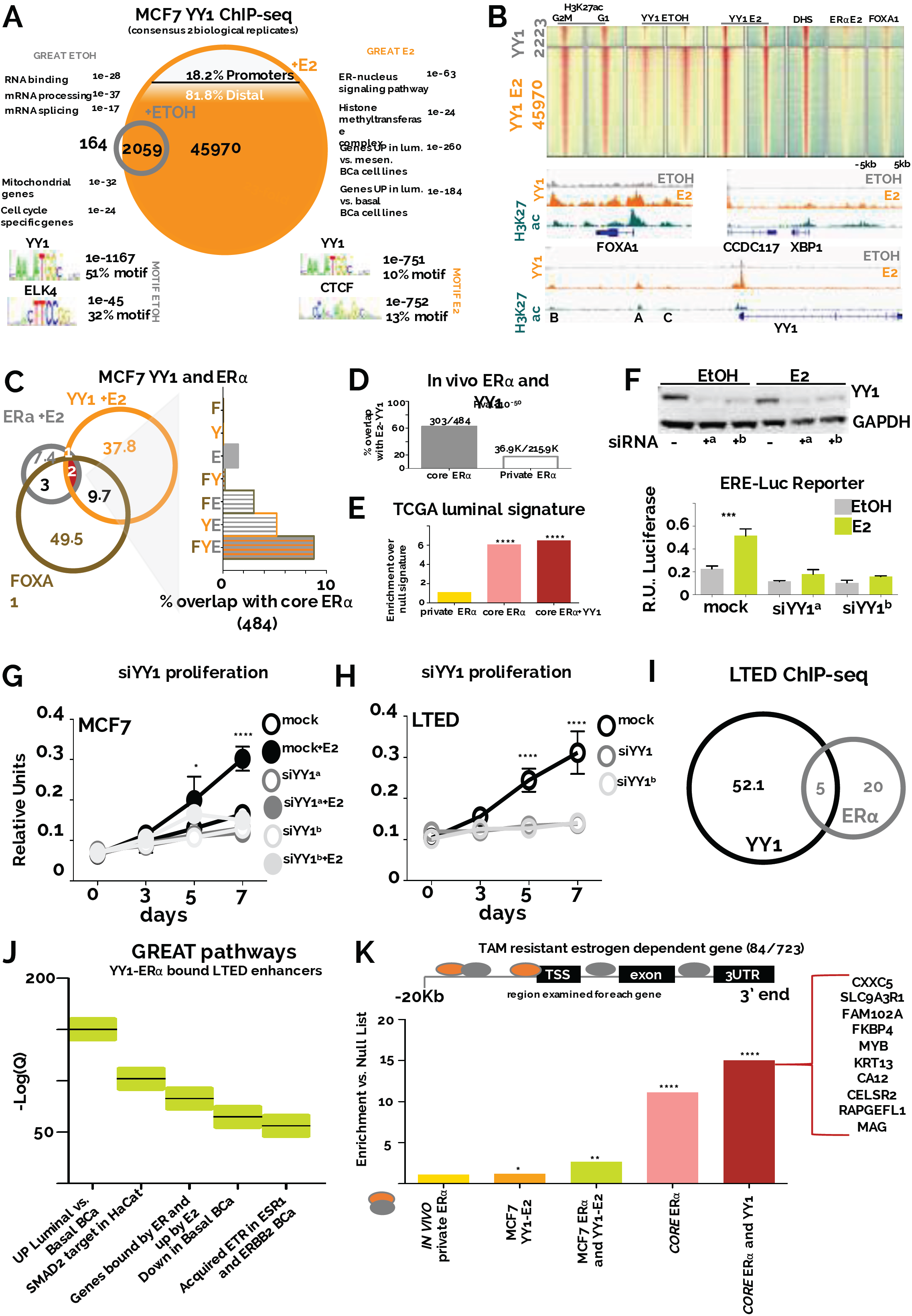
YY1 marks critical enhancers in breast cancer cells. A) ChIP-seq data from ER-positive MCF7 for YY1 in quiescent or 17ß-estradiol (E2) stimulated cells B) Heatmaps showing global enrichment profiles of several chromatin markers associated with active regulatory regions in MCF7 cells C) Overlap between ER, YY1 and FOXA1 in MCF7 cells. The right panel shows the potential overlap with *in vivo-* derived core ERα binding sites D) ERα core binding sites are strongly enriched for YY1 binding in MCF7 cells while patient-specific ERα bindings are generally YY1-free. E) Genes used to classify luminal breast cancer patients are strongly enriched for ER-YY1 binding sites. Asterisks represent p<10^−5^ in a Fisher’s Exact test vs. private ER F) YY1 depletion leads to transcriptional shut-down of an ERE-driven luciferase reporter. Bars and error bars represent the average of 5 independent experiments with SE. Asterisks represent significance at P<0.001 after ANOVA with Dunnet’s correction. G) Silencing YY1 blocks estrogen-induced growth in MCF7 cells H) YY1 depletion leads to growth arrest in AI resistant LTED cells. Proliferation assays were conducted in biological triplicate. Error bars indicate 95% confidence intervals. Asterisks represent significance at P<0.05, 0.01, 0.001 and 0.0001 after 2-way ANOVA with Tukey’s post-test I) Overlap of YY1 and ERα binding sites in LTED cell lines J) ERα-YY1 bound enhancers in LTED cells underlie the transcription of genes associated with luminal breast cancer and acquired endocrine therapy resistance K) core ERα-YY1 bound enhancers are strongly enriched near estrogen responsive genes that are not suppressed by Tamoxifen co-treatment.

Our *in vivo* analysis suggest that clonal YY1-bound enhancers are generally not enriched for EREs or ERα, with the exception of the atypical core-ERα (Fig 2B). In agreement, our *in vitro* data show only marginal overlap between YY1 and ERα or its pioneer factor FOXA1 (Fig. 4B-D) indicating that generally YY1 recruitment is independent from ERα. On the other hand, YY1, ERα and FOXA1 co-localize at increased frequencies at core-ERα loci in MCF7 cells (Fig. 4C). Similar observations were made by comparing YY1 overlap with *in vivo* derived ERα binding (60% overlap with core ERα vs. 18% overlap with patient-unique ERα). In addition, we find that genes defining the luminal subtype in TCGA patients are significantly enriched for ERα core binding with YY1 but not patient-unique ERα (Fig. 4D). Overall, these data further suggest that YY1 might stabilize ERα binding^42^ at a small subset of transcriptionally productive enhancers (core-ERα) captured in most tumor cells and most patients^43^. To test if YY1 can contribute to ERα driven transcription, we measured luciferase activity from a promoter driven by an array of estrogen response elements (EREs) in MCF7 cells in presence or absence of YY1(Fig. 4E) and show absolute dependencies on YY1. Furthermore, YY1 depletion also abrogates cell proliferation in response to estrogen stimulation in MCF7 (Fig. 4F) suggesting that YY1 is a direct driver of the clonal proliferation observed in the BCa (Fig. 3D-E). These observations were replicated in other independent luminal BC cell models (ZR75 and T47D, Supplementary Figure 8A-B). Finally, we show that YY1 depletion leads to significant downregulation of core-ERα target genes in luminal BC cell line models (Supplementary Figure 8C). Collectively these data identify YY1 as a novel essential transcription factor significantly contributing to ERα regulatory network transcriptional activity.

### YY1 contributes to drug-resistance in luminal BC

YY1 motif is highly enriched in clonal enhancers identified in primary and metastatic luminal patients (Fig 2B). All metastatic patients included in this study relapsed following adjuvant endocrine therapies suggesting that YY1 might also play role in this setting. In agreement, primary and metastatic samples show clonal YY1 enhancer activity, indicating that YY1 positive cells are not effectively cleared by the therapy (Fig. 3A). Therefore, we investigated the role of YY1 in LTED cells, an MCF7-derivative that develop estrogen-independent growth partly through constitutive activation of ERα signalling^16^. YY1 depletion leads to complete abrogation of LTED growth demonstrating that YY1 is still required at this stage (Fig. 4G). Interestingly, LTED cells have an expanded repertoire of ERα binding compared to MCF7, fuelled by endogenous ligands ^13,16^. The set of enhancers engaged by ERα and YY1 in LTED cells is radically different compared to MCF7, with the majority of ERα-YY1 being specific to each cell type (Fig 4I, LTED only: 3598/5037). ERα-YY1 bound enhancers in LTED strongly associates with the transcription of genes involved with acquired endocrine therapy, suggesting that during epigenetic reprogramming^16^, YY1 might stabilize ERα to LTED specific enhancers (Fig. 4J). To further examine the relationship between YY1 and endocrine resistance we analyzed a set of estrogen responsive genes whose transcription cannot be antagonized by Tamoxifen treatment in MCF7 cells^58^. These genes were not enriched for patient-private ERα, but we saw an ever-increasing association with ERα-YY1 bound enhancers, especially with core ERα-YY1 (Fig. 4K). For example, ERα-YY1 is found near CXXC5 and SLC9A3R1, ranked respectively first and second as the most strongly estrogen-induced gene that cannot be antagonized by Tamoxifen ^58^. Collectively, these data strongly support the role of YY1 in ERα BC growth and progression.

### YY1-ERα promote SCL9A3R1 expression despite endocrine treatment

SLC9A3R1 (NHERF1/EBP50)^59^ SLC9A3R1 encodes a Na/H exchanger regulatory cofactor. SLC9A3R1 null mice have disrupted protein-kinase-A-dependent cAMP-mediated phosphorylation^60^. In agreement with SLC9A3R1 potential role in endocrine resistance, meta-analysis of patient-derived data using all available genes (n=22,277) reveals that SLC9A3R1 expression is amongst the top 1% of genes with the strongest prognostic association with relapse in a cohort of 724 ET-treated ERα-positive patients ^39^ (Fig. 5A). High expression of SLC9A3R1 also significantly correlates with poor survival in additional independent ERα-BC datasets (Supplementary Figure 9A). In addition to Tamoxifen treatment, SLC9A3R1 remains transcriptionally active in most endocrine therapy resistant BC cell lines that retain ERα expression (Supplementary Figure 9B) but genetic or pharmacological (Fulvestrant) suppression of ERα is sufficient to block SLC9A3R1 transcription (Supplementary Figure 9B-C). Specifically, SLC9A3R1 expression is not antagonized by estrogen deprivation^61^ (LTED models, Supplementary Figure 9B-D), nor Raloxifene *in vitro* (Supplementary Figure 9E) or neo-adjuvant AI treatment in clinically treated patients *in vivo* (Fig. 5B). Overall, these data demonstrate that SLC9A3R1 is a direct ERα target whose expression cannot be antagonized by first-line endocrine therapies.

**Figure 5:**
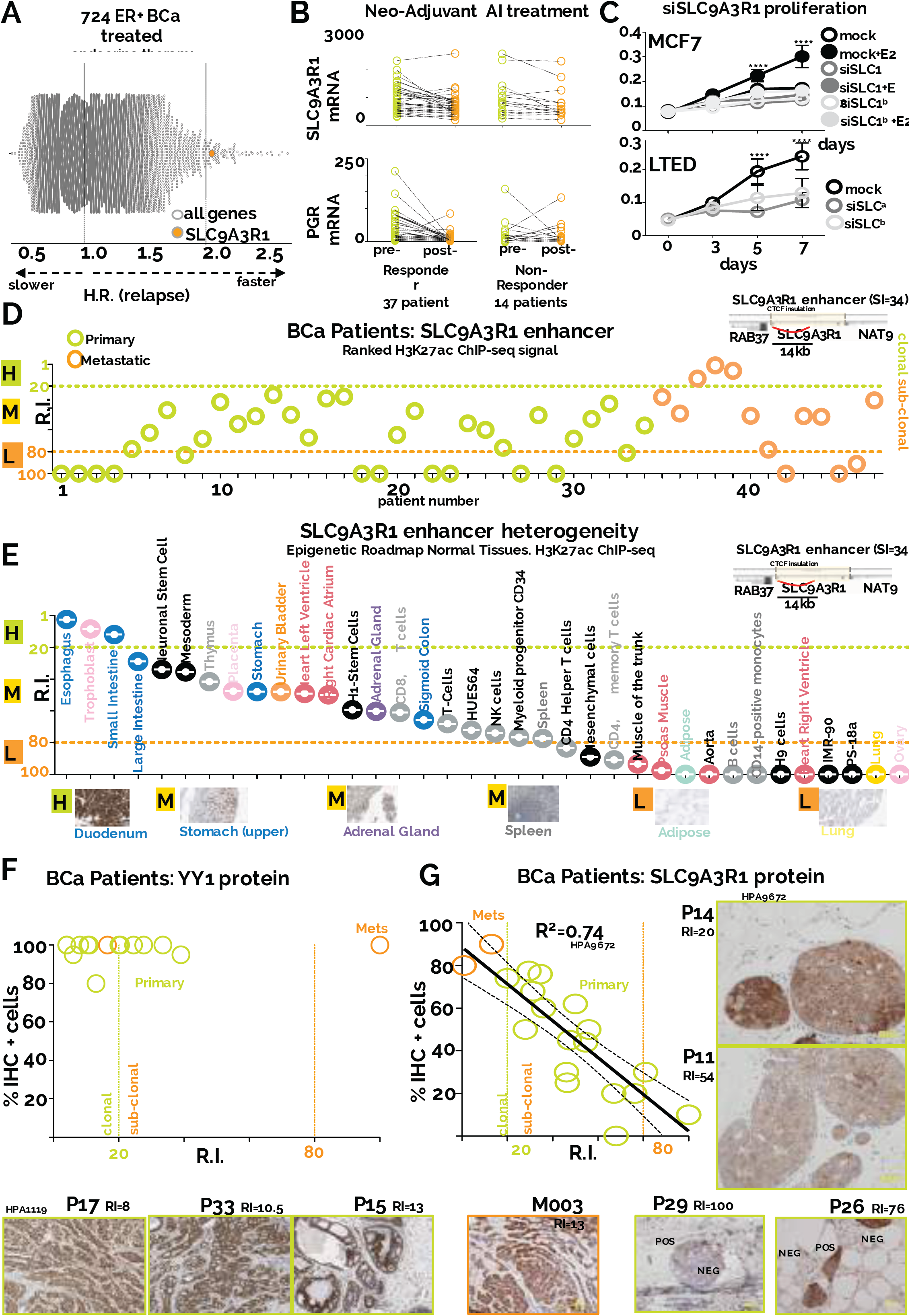
Epigenomic mapping predicts the size of phenotypic clones in patients. A) Global Kaplan-Meier analysis summarize univariate analysis for each gene included in the Affymetrix microarray platform. Hazard Ratios are plotted in the X axis B) SLC9A3R1 RNA levels pre- and post-short-term aromatase inhibitor treatment in responder and non-responder patients^61^. Oestrogen-dependent expression of progesterone receptor mRNA is shown as comparison C) Silencing SLC9A3R1 leads to proliferation arrest in response to estrogen stimulation in MCF7 and estrogen independent growth in LTED cells. Proliferation assays were conducted in biological triplicate. Error bars indicate 95% confidence intervals. Asterisks represent significance at P<0.05, 0.01, 0.001 and 0.0001 after 2-way ANOVA with Tukey’s post-test D) RIs for the SLC9A3R1 enhancer within all the individual patients included in the current study. SLC9A3R1 enhancer location and its 3D interactions are shown in the top right inset E) SLC9A3R1 enhancer ranking analysis of available Epigenome Roadmap H3K27ac datasets. Tissues are displayed from the strongest to the weakest SLC9A3R1 enhancer activity (based on RI). Representative IHC analysis of normal tissues stained with a SLC9A3R1 antibody are shown. F-G) YY1 and SLC9A3R1 IHC analysis of BC patients profiled using H3K27ac ChIP-seq. Predicted activity (RI) of YY and SLC9A3R1 enhancers is shown on the X axis. The number of cells positively stained for YY1 and SLC9A3R1 protein is indicated on the Y axis. Linear regression R square, confidence intervals and representative staining are also shown.

Bulk RNA-seq data from a panel of cancer cell lines demonstrate that ERα-positive BC cells have the highest levels of SLC9A3R1 mRNA (Supplementary Figure 10A). More importantly, TCGA RNA-seq analysis shows that SLC9A3R1 expression is higher specifically in ERα BC patients compared to normal tissue or other subtypes (Supplementary Figure 10B). Chromatin analyses of MCF7 and LTED cells identify ER-bound enhancers at 3 independent loci within the insulated SLC9A3R1 locus (E1-E3), a RD region directly looping to the YY1-bound SLC9A3R1 promoter within a CTCF insulated perimeter (Supplementary Figure 10C). Strikingly, E1 and E2 contain core ERα bindings in addition to YY1, while E3 contains a patient unique ERα binding with no YY1 (Supplementary Figure 10C). *In vivo* transcriptional analysis demonstrates that SLC9A3R1 is the only gene near the E1-E2 enhancers that shows a significant increase in bulk-RNA level when comparing normal breast tissue with ERα–positive BC (Supplementary Figure 10D). Interestingly, enhancer-activity appears to be immune to endocrine therapy (MCF7 vs. MCF7 tamoxifen resistant and LTED, Supplementary Figure 10C). Collectively, these data strongly support the notion that SLC9A3R1 expression is driven by a breast cancer specific enhancer within the expanding ERα-YY1 clone during tumor initiation. Nonetheless, SLC9A3R1 expression is dependent on YY1 (Supplementary Figure 11A), demonstrating that both ERα and YY1 are essential for full enhancer activity. Silencing SLC9A3R1 is sufficient to abrogate oestrogen-induced growth in ERα-positive cells (Fig 5C). Intriguingly, SLC9A3R1 is not essential for a second in ERα-positive model (T47D) but appears to be a critical gene for both AI-resistant cells models (Fig. 5C and Supplementary Figure S11B). Collectively, these data demonstrate that ERα-YY1 regulate SLC9A3R1 via enhancer binding and identify SLC9A3R1 as a novel player involved in ET resistance.

### Mapping phenotypic heterogeneity using YY1 and SLC9AR1 enhancer activity

Both SLC9A3R1 and YY1 enhancers are commonly activated in our patient’s dataset (SI=34 and SI=41 respectively). Yet, YY1 enhancer identifies YY1-positive cells as a dominant clone in almost all patients (RI≤20, Fig 3A). Conversely, SLC9A3R1 enhancer activity indicates that SLC9A3R1 marks a potentially dynamic sub-clonal population in most primary patients (RI≥20, Fig 5D). Our *in vitro* data suggest that SLC9A3R1 transcription cannot be antagonized by endocrine therapies while SLC9A3R1 is important for resistant BC cell lines. This predicts that the SLC9A3R1-positive population should increase under adjuvant treatment. Interestingly, the only evidence of a clonal SLC9A3R1 population was found in samples from three metastatic, endocrine-resistant patients (Fig. 5D).

Bulk transcriptional data show that average SLC9A3R1 expression is significantly higher in ERα positive BC cells but do not inform about potential subpopulations (Supplementary Figure 10A-B). Conversely, meta-analysis of SLC9A3R1 enhancer activity (RI) within the ENCODE H3K27ac datasets indicates that MCF7 cells contain a clonal SLC9A3R1 population while all other cell lines appear to have decreasing sub-clonal populations (Supplementary Figure 11C). Of note, the size of the sub-clonal population correlates with total RNA content for the cells contained in both assays, suggesting that the decreasing bulk RNA signal is driven by a progressively smaller subpopulation (Supplementary Figure 11C). Similar analysis of YY1 enhancer indicate that cancer cell lines are vastly clonal for YY1 expression (Supplementary Figure 11D) in agreement with clinical samples. Notably, both YY1 and SLC9A3R1 RIs in mammary epithelial cells suggest the presence of sub-clonal populations. These observations fit well with experimental data from IHC profiles from normal breast (Fig. 3D and Supplementary Figure 12B). Meta-analyses of H3K27ac from the Epigenome Roadmap database predict that most tissues potentially have only sub-clonal populations, as determined by SLC9A3R1 enhancer activity (Fig. 5E). In agreement, the size of the sub-clonal population tracks the mRNA signature of each tissue with SLC9A3R1 potentially clonal tissues accumulating the highest amount of RNA-seq tags (Small and Large Intestine, Supplementary Figure 12). Analogously to YY1, meta-analysis of IHC data identifies decreasing SLC9A3R1-positive with increasing RI scores (Fig. 5E and Supplementary Figure 12B). Finally, to further validate that RI index can estimate phenotypic clones, we retrospectively collected available FFPE biopsies for the BC patients profiled with H3K27ac ChIP-seq (n=19). We then performed IHC using YY1 (Fig. 5F) and SLC9A3R1 (Fig. 5G) antibodies and compared the predicted enhancer activity (RI) with the actual size of YY1 and SLC9A3R1-positive clones within each patient. With the exception of one metastatic sample (M3), YY1 staining robustly correlate with RI, confirming large clonal YY1 positive populations in all examined tissues (Fig. 5F). In parallel, SLC9A3R1 enhancer activity correctly estimated the size of the sub-clonal subpopulations in individual patients (Fig 5D). Further meta-analyses on Protein Atlas data support these findings, by identifying YY1 clonal populations and SLC9A3R1 sub-clonal populations in most ERα BC samples (Supplementary Figure 13A-C). Overall, these data strongly support the notion that enhancer activity can be used to qualitatively deconvolute heterogeneous populations into phenotypical subclones.

### Phenotypic evolution during BC progression is shaped by endocrine treatment

Tumor evolution studies have primarily focused on treatment naïve patients, taking advantage of multi-regional sampling to retrospectively monitor changes in clonality^14,62^. Clonal tracking is dependent in part on passenger mutations, and the effect of therapy has been rarely accounted for^13,63^. More importantly, clonality has been traced uniquely using genetic variants, with the intrinsic limitation of correlating genetic changes to phenotypic ones. For example, sub-clones defined by passenger mutations might be phenotypically equivalent, while a recent study using barcoded glioblastoma cells shows that phenotypic clones might evolve independently from mutational signatures ^2^. In addition, the few studies that looked at driver mutations in coding regions of primary and metastatic BC disease found relatively similar mutational landscapes^15^, suggesting that mapping phenotypic clones though BC progression might reveal new targets. Our ability to acquire qualitative estimates of phenotypic clones using enhancer ranking provides for a potential approach for tracking changes in tumor heterogeneity with the additional advantage of predicting for potentially functional changes. We interrogated our patient’s dataset focusing on events occurring between treatment-naïve primaries and treatment-resistant metastatic BC (Fig. 6A). We hypothesized that phenotypic clonal evolution might be driven by a coordinated activation/selection of groups of enhancers during BC progression and this could be influenced by treatment. Our previous results suggest that YY1+ cells remain clonal during progression (Fig 3A). Conversely, we show that SLC9A3R1 expression is not antagonized by endocrine treatment suggesting that SLC9A3R1-positive clones could expand during progression. We then calculated changes in RI (ΔRI) for all enhancers captured in at least three patients (SI>3, n=88935) between primary and metastatic samples (Fig. 6B). SLC9A3R1 ranks amongst the enhancers with the strongest increase in predicted clonality going from primary to metastatic samples (Fig. 6B-C, 3.86σ from median ΔRI). Conversely, YY1 enhancer activity remains relatively unchanged (Fig. 6B-C). These data support our initial hypothesis and suggest that SLC9A3R1-positive clones might expand in response to treatment. To substantiate these data, we mapped the size of YY1 and SLC9A3R1 phenotypic clones using IHC in an independent series of 20 matched longitudinal biopsies. All surgical biopsies were obtained from treatment naïve patients, while all the metastatic biopsies were taken at first relapse after endocrine treatment^13^. We found YY1+ cells clonal both in primary and metastatic biopsies (Fig. 6D). Conversely, SLC9A3R1+ subclones significantly expand during metastatic progression to become completely clonal (100% staining) in 13/20 patients. Interestingly, the only metastatic case in which we have observed a contraction of the SLC9A3R1+ clone also showed a concomitant loss of ERα and PR positivity, confirming our *in vitro* analysis and demonstrating that SLC9A3R1 remains an ERα dependent-target despite being ET insensitive *in vivo* (Fig. 6D, red line and Supplementary Figure 9B-E). Overall, these data demonstrate that changes in enhancer ranking can estimate functional evolution during breast cancer progression.

**Figure 6:**
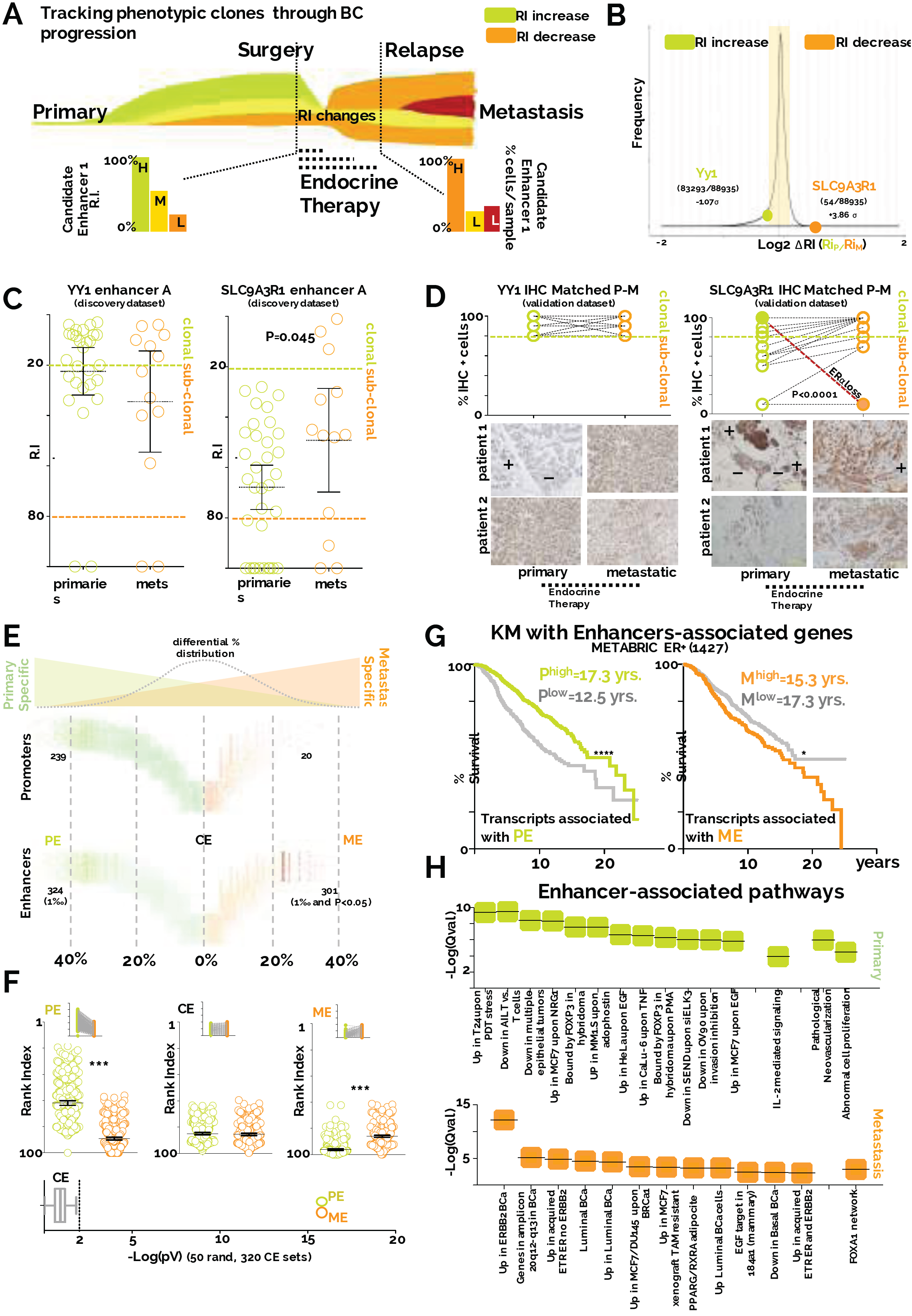
Endocrine treatment shapes phenotypic evolution. A) Theoretical framework of the analysis. The relative size of phenotypic clones can be tracked using enhancer activity (RIs). Phenotypic clones can be positively or negatively selected during BC progression in response to endocrine therapies. B) Expanding or contracting phenotypic clones were defined based on the RI-ratio in primary and metastatic samples (RI_P_/RI_M_). Distribution of RI-ratio identified top candidate enhancers YY1 RI does not change significantly during progression, while SLC9A3R1 RI ranks among the enhancers with stronger increase in activity during progression. Vertical bars represent σ (Standard Deviation) increments from the population median C) Scatterplot of YY1 and SLC9A3R1 enhancer ranking according to patient stage. Bars indicate mean and 95% confidence intervals. Asterisks represent significance at P<0.05 after students two-tail T-Test D) IHC staining for YY1 and SLC9A3R1 positive cells in an independent matched longitudinal cohort of ERα breast cancer patients. All normal and primaries are treatment naïve. All metastatic have received endocrine therapies (Tamoxifen or Aromatase inhibitors). Statistical significance was calculated using a pair-wise, two-tail T-test. Representative images are also shown E) Enhancer and promoter stratification based on frequency of usage in primary and metastatic patients. Percentages were calculated for each regulatory region for each stage (primary and metastatic) and differential was then derived and plotted on the X-axis F) RI indexes for all PE and ME are plotted. As a control, RI for common enhancer (CE) are also plotted. Permutation was used to assess changes in RI in 50 randomly selected sets of CE G) Kaplan-Meier analysis using averaged RNA expression of genes associated with PE or ME regulatory regions. Genes were assigned considering CTCF insulated perimeters E) Pathway analysis for genes associated with PE or ME regulatory regions. Pathways were identified using GREAT and are listed in order of significance (qValue).

To gain more insight on functional evolution, we systematically annotated all regulatory regions based on bias in detection between primary and metastatic patients (Fig 6E). As expected, the bulk of enhancers and promoters do not show bias toward primary and metastatic BC patients (common enhancers, CE). However, we could successfully identify two distinct sets of regulatory regions that are preferentially associated with primary (primary enhancers, PE) or metastatic (metastatic enhancers, ME) patients. Remarkably, while CE do not show stage-specific changes in RI, PE underlie larger sub-clonal populations in primary cancers (statistically higher RIs in primary compared to metastatic, Fig. 6E). Likewise, ME have lower RI in metastatic samples suggesting that the number of cells carrying these enhancers have increased during progression (Fig. 6E). We next explored the potential causes and functional consequences driving these coordinated epigenetic changes. We thus identified the potential transcriptional targets of our enhancers taking in account CTCF boundaries^37^. Strikingly, we find that PE-associated gene-transcription is associated with significantly better outcome while ME-associated gene-transcription in primary samples is associated with poor prognosis (Fig 6D). These data imply that primary samples containing larger subpopulations of phenotypic clones with metastatic features relapse earlier.

We then mined PE and ME regulatory regions to identify the associated biological features^56^. PE appear to promote abnormal proliferation and vascularization, two key events in early tumorigenesis. Remarkably, metastatic samples switch to functional clones characterized by genes associated with BC progression (FOXA1^43^) or endocrine therapy resistance^64,65^ (Fig. 6E). Altogether, these data suggest that endocrine therapies play a central role in shaping phenotypic clonal evolution. Additional in-depth studies are needed to dissect the temporal events triggered during phenotypic clonal evolution. Phenotypic subclones could evolve by early coordinated activation and decommissioning of epigenetically defined regulatory regions (*acquired*), selection of the fittest pre-existent epigenomic landscape (*de novo*) or a combination of both.

### Discussion

Our work describes the first systematic epigenetic profiling of primary and metastatic luminal breast cancer and reveals several critical principles underlying phenotypic-functional heterogeneity and its role in breast cancer progression. By mapping H3K27ac in untreated and treated patient samples we have also identified YY1 and SLC9A3R1, two new key players contributing to BC. While genomic profiling of breast cancer patients has revealed extensive clonal heterogeneity and retrospective tumour evolution^28,66^, the vast majority of the mutational burden can be considered composed of passenger mutations^29,67^ making difficult to extrapolate actual phenotypes. Most RNA-based analysis, which may better reflect the phenotypic state of cancer cells, is generally obtained from bulk tissue and cannot inform on the existence of distinct subpopulations. Finally, molecular pathology can inform on the relative amount of protein abundance at the single-cell level but is laborious and not suitable for testing multiple targets simultaneously. In this work, we used epigenomic analyses to extrapolate phenotypic heterogeneity in solid tumour samples. Our analysis reveals that histone-based ChIP-seq signals, similarly to ATAC-seq^32^, generally correlates with the number of cells in a population carrying the specific epigenetic information. Our predictions using YY1 and SLC9A3R1 enhancer fit extremely well with experimental data derived from normal tissues or BC patients. The findings that clonal regulatory regions dominating the landscape of individual tumor samples are shared across many patients, parallel recent genomic evidences showing that truncal (high allele frequency) mutations are also the most common mutations within cancer cohorts.

The results described here have several practical implications for BC. First, by comparing samples from drug-resistant metastatic patients with drug-naïve primary samples, we uncovered a set of enhancers marking phenotypic clones that significantly expand during breast cancer progression. Notably, these enhancers are strongly associated with genes specifically transcribed in cells that acquire endocrine therapy resistance (Fig. 6H). Conversely, enhancers progressively lost during tumour progression are linked to processes that often occur early in tumorigenesis. A set of enhancers expanding in metastatic samples point at progressive activation of FOXA1 and its network. It was recently reported that FOXA1 levels are increased in metastatic samples^43,68^. Our data then predict that, similarly to SLC9A3R1, FOXA1 positivity increases as a consequence of the expansion of a phenotypic clone marked by an active FOXA1 enhancer and not via increased transcription of the FOXA1 gene within single cells. It is tempting to speculate that this paradigm might be valid for other genes. If correct it might signify that during cancer evolution, the proportion of cells activating transcription is more important than the absolute changes in transcription at the single cell levels. Interestingly, a set of enhancers deactivated during progression involve IL-2 signalling (Fig. 6H). Reduction in IL-2 signalling was identified as a potential marker of relapse^69^. Whether the IL-2 signal source is the BC cells^70^ themselves or it is due to a small contamination of immune cells, needs to be defined. Equally, it will be important to measure real-time activation/selection of enhancers in appropriate systems to ultimately establish if phenotypic cancer evolution can be driven by Lamarckian events.

Finally, our analysis has identified two novel drivers of luminal BC. Firstly, we identified YY1 as a key TF associated with clonal enhancers and promoters in BC patients. Our data strongly support the idea that YY1 acts as a global co-activator in cancer cells associating with the entire active epigenetic landscape. Several lines of evidence indicate that YY1 might interact directly with modified nucleosomes, possibly through its partner INO80^57^. YY1 widespread association with clonal enhancer suggests it might play a role in epigenetic memory. Intriguingly, a positive screen for factors that improve induced pluripotent cells formation (iPS), identified YY1 as the top hit, further supporting its potential role as enhancer gatekeeper ^71^. More specifically to ERα BC, we hypothesize that YY1 plays a critical role to stabilize ERα binding at the transcriptionally productive core- ERα enhancers. Single-molecule imaging shows that estrogen activated ERα increases its residency time on the chromatin^42^ and recent evidence has shown that eRNA can trap YY1 on the chromatin ^49^. More importantly, enhancer co-occupancy for YY1 and ERα occurs almost exclusively at highly shared-highly functional core-ERα bound loci. Altogether, these data raise the intriguing hypothesis that YY1 might contribute to increased ERα residency at clonal enhancers. This could explain why some ERα are captured in most patients, as longer residency time would increase chances of being captured by ChIP-Seq^43^. Longer residency might also explain the increased transcriptional activity (Fig. 4D) and increased TF footprints (Fig. 2C) of these enhancers.

YY1-ERα jointly control SLC9A3R1 enhancer activity, an event that cannot be antagonized by conventional first-line endocrine therapies (Tamoxifen or AI) and that drives SLC9A3R1 clonal expansion during breast cancer progression. Of note, primary patients with high SLC9A3R1 expression might be viewed as containing larger population of SLC9A3R1+ cells, thus resembling drug-resistant BC. Intriguingly, SLC9A3R1 is amongst the strongest single prognostic genes for relapse-free survival when considering endocrine treated patients (Fig. 5A). An attractive possibility is that YY1 stabilizes ERα sufficiently at the SLC9A3R1 enhancer maintaining epigenetic memory in the presence of external antagonists. Future studies are required to investigate the exact mechanisms through which SLC9A3R1 contribute to BC and efficient strategies to antagonize its transcription, possibly using CDK4/6 inhibitors^72^ to destabilize YY1. We recently demonstrated that individual endocrine therapies can drive parallel genetic evolution *in vivo* ^13^ and epigenetic reprogramming *in vitro*^16^. Our data now strongly support the notion that therapeutic interventions also play an essential role driving specific epigenetic evolution during BC progression in the clinic. Metastatic re-biopsy at the time of relapse, which is becoming commonplace in clinical practice should then examine epigenetic changes in addition to newly acquired genomic ones, especially when no new genetic drivers to guide further treatment are apparent^15^

## On Line Materials and Methods

### Tumour tissue processing

Breast cancer sample for ChIP-seq were collected by Imperial Tissue Bank (project ethic approval R15021) and from Breast Cancer Now Tissue Bank (BCNTB-TR000053-MTA & TR000040). Breast cancer fresh frozen tissue samples each undergo aseptic macroscopic adipose tissue dissection. The dissected tumour tissue is sectioned into 2mm × 2mm fragments in a petridish placed over dry ice. Tumour fragments are then fixed using 1% formaldehyde solution for 10 minutes. Cold glycine (1M) is added to the formaldehyde-fixed tissue for 10 minutes. The tumour fragments are then pulverised using pestle and mortar and homogenised using liquid nitrogen.

### Chromatin immunoprecipitation (ChIP)

The ChIP protocol was conducted as described by Schmidt et al.^73^ with few modifications. In summary, following fixation, the tumour tissue undergoes chromatin extraction and sonication using the Bioruptor Pico sonication device (Diagenode; B01060001) using 20 cycles (30s on and 30s off) at maximum intensity. Purified chromatin was then separated for 1. Immunoprecipitation using 4ug of H3k27ac antibodies (Abcam; ab4729) per ChIP experiment or using 4ug of YY1 antibodies (Santa Cruz; sc-281 X). ChIP-seq experiment for YY1 were performed in biological duplicates. 2. Non-immunoprecipitated chromatin, used as Input control and 3. Assessment of sonication efficiencies using a 1% agarose gel. Before construction of ChIP-seq libraries (NEB Ultra II kit, see supplementary methods), enrichment of the immunoprecipitated sample was ascertained using positive and negative controls for ChIP-qPCR. Library preparation was performed using 10 – 50 ng of immunoprecipitated and Input samples.

### ChIP-qPCR

Briefly, reactions were carried out in 10 ul volume containing 5 ul of Sybergreen mix (ABI; 4472918), 0.5 ul of primer (5 uM final concentration), 2.5 ul of genomic DNA and 2 ul of DNASE/RNASE –free water. A three-step cycle programme and a melting analysis were applied. The cycling steps were as follows: 10s at 95 oC, 30s at 60 oC and 30s at 72 oC, repeated 40 times.

### Ranking and Sharing Index

See Supplementary Computational Methods.

### VSE

See Supplementary Computational Methods.

### DHS imputations and TF motif analyses

See Supplementary Computational Methods.

### Imputed DHS with vivo ERα binding Overlap

ERα binding from in breast cancer patients were obtained from ^43^. ERα sharing index was calculated as before (see Supplementary Computational Methods). Overlap with imputed DHS was calculated using BedTools calculating the overlap (at least one base pair) via Cistrome Pipeline Analysis Suite (http://cistrome.org/Cistrome/Cistrome_Project.html). The percentage of overlap were calculated using binned DHS as variable first dataset and all the concatenated in vivo ERα as second dataset.

### Footprint analysis

See Supplementary Computational Methods.

### Encode and Epigenomic Roadmap Ranking

See Supplementary Computational Methods.

### Immunocitochemistry

Hematoxylin and eosin staining of clinical samples was performed to calculate tumor burden prior to ChIP-seq. Briefly, 4-μm-thick sections were obtained from formalin-fixed and paraffin-embedded specimens. After de-waxing in xylene and graded ethanol, sections were incubated in 3% H2O2 solution for 25 minutes to block endogenous peroxidase activities and then subjected to microwaving in EDTA buffer for antigen retrieval. For YY1 (Protein Atlas HPA001119, <u>Atlas Antibodies</u> <u>Cat#HPA001119, RRID:AB_1858930</u>) the flowing conditions were used: tissue sections were incubated with the primary monoclonal. overnight at 4°C, and chromogen development was performed using the Envision system (DAKO Corporation, Glostrup, Denmark). A minimum of 500 tumor cells were scored with the percentage of tumor cell nuclei in each category recorded. For SLC9A3R1 (HPA9672 and HPA27247, <u>Atlas Antibodies Cat#HPA009672, RRID:AB_1857215</u> and <u>Atlas Antibodies Cat#HPA027247, RRID:AB_10601162</u> respectively) the following conditions were used. HPA9672 was diluted 1:400 and HPA27247 was diluted 1:1500. Staining was automatized with a Ventana Benchmark Ultra using epitope retrieval ER2 for 20 minutes. ER and PgR immunoreactivity was assessed by the FDA-approved ER/PR PharmDX kit (Dako). The prevalence of ER/PgR positive invasive cancer cells, independent of their staining intensity, was quantitatively annotated in the original reports. In accordance with ASCO/CAP guidelines, tumours with ≥1%of immunoreactivity were considered positive

### Cell culture

MCF7 was cultured using Dulbecco’s modified Eagle’s medium (DMEM) containing 10% fetal calf serum (FCS) and 100 U penicillin/0.1 mg ml^−1^ streptomycin, 2mM L-glutamine plus 10^−8^ 17-β-estradiol (SIGMA E8875). MCF7 long term oestrogen deprived (MCF7-LTED) cells were grown in phenol-free DMEM with 10% charcoal-stripped FCS (DCFCS) and 100 U penicillin/0.1 mg ml^−1^ streptomycin and 2mM L-glutamine. T47D and T47D-LTED cells were passaged using DMEM containing 10% FCS and 100 U penicillin/0.1 mg ml^−1^ streptomycin, 2mM L-glutamine and phenol-free DMEM with 10% DCFCS and 100 U penicillin/0.1 mg ml^−1^ streptomycin and 2mM L-glutamine, respectively. ZR75-1 cells were grown in DMEM containing 10% FCS and 100 U penicillin/0.1 mg ml^−1^ streptomycin, 2mM L-glutamine.

### sIRNA

Small interfering RNA (siRNA) against SLC9A3R1 (Gene ID; 9368: Ambion; s17919, s17920), YY1 (Gene ID; 7528: Ambion; s14958, s14959, s14960) and *Silencer* negative control (Ambion; AM4611). 1.5 × 10^5^ cells were seeded per well using a 6-well plate. MCF7 cells were seeded in phenol-free DMEM with 10% DCFCS and 100 U penicillin/0.1 mg ml^−1^ streptomycin and 2mM L-glutamine. Following 24 hours, the cells were then transfected with siRNA using Lipofectamine 3000 (Invitrogen; L3000015). T47D and ZR75-1 cells were seeded in DMEM containing 10% FCS and 100 U penicillin/0.1 mg ml^−1^ streptomycin, 2mM L-glutamine. Following 24 hours, the cells were then transfected with siRNA using Lipofectamine 3000 (Invitrogen; L3000015). Cells were harvested for analysis following at least 48 hours of transfection.

### Cell lysis and Western blot

Cells were washed twice in ice-cold PBS and lysed in RIPA (Sigma-Aldrich; R02780) buffer supplemented with protease (Roche 11697498001) and phosphastase (Sigma-Aldrich 93482) inhibitors for 30 minutes with intermittent vortexing. Samples were centrifuged at 4^o^C at maximum speed for 30 minutes after which, the supernatant is transferred to a clean Eppendorf. Protein concentrations for each sample was ascertained using the bicinchoninic acid (BCA) assay (ThermoFisher Scientific; 23227). Equal amounts of lysates were loaded into BOLT 4-12% Bis-Tris Plus Gel (Invitrogen; NW04120BOX). Proteins were transferred to a Biotrace nitrocellulose membrane (VWR; PN66485) and incubated with primary antibodies overnight. Proteins were then visualised using goat anti-mouse (ThermoFisher Scientific; 31446) and anti-rabbit (ThermoFisher Scientific; 31462) HRP conjugated secondary antibodies. Amersham ECL start Western Blotting Detection reagent (GE Healthcare Life Sciences; RPN3243) was used for chemiluminescent imaging using the Fusion solo (Vilber; Germany) imager.

### Transcriptional profiling

Following 48 hours of transfection, MCF7 cells were either treated with 10^−8^ 17-β-estradiol (SIGMA E8875) or control treatment for 6 hours prior to RNA extraction. T47D and ZR75-1 cells lines were harvested for RNA following 48 hours of transfection. No treatments were added.

### RNA extraction and real-time PCR

Total RNA was extracted using RNeasy Mini Kit (Qiagen; 74106), and the cDNA was reverse transcribed from 1ug of RNA using iScript cDNA synthesis kit (Bio-Rad; #1708891). Real time-qPCR (RT-qPCR) reactions were carried out in 10 uL volume containing 5 uL of sybergreen mix (ABI; 4472918), 0.5 ul of primer (2.5 uM final concentration), 2.5 ul of genomic DNA and 2 ul of DNASE/RNASE–free water. A three-step cycle programme and a melting analysis were applied. The cycling steps were as follows: 10s at 95 ^o^C, 30s at 60 ^o^C and 30s at 72 ^o^C, repeated 40 times^19^.

### Luciferase reporter assay

MCF7 cells were seeded in a 24-well plates at 5 × 10^4^ cells per well in phenol-free DMEM with 10% DCFCS and 100 U penicillin/0.1 mg ml^−1^ streptomycin and 2mM L-glutamine. After 24 hours of incubation, transfection of plasmid DNA was performed using Lipofectamine 3000 (Invitrogen; L3000015). Cells were transfected with 100ng of ERE_Luciferase reporter, 10ng of the renilla luciferase control plasmid (pRL-CMV), 10ng of pSG5_ER-α, 15 nm of siRNA and 280ng of Bluescribe DNA (BSM) per well; totalling 400ng of DNA/well. After 12 hours of transfection the media was replaced with fresh phenol-free DMEM with 10% DCFCS and 100 U penicillin/0.1 mg ml^−1^ streptomycin and 2mM L-glutamine. Treatment with 10^−8^ 17-β-estradiol (SIGMA E8875) or control treatment was administered and the cells incubated for 24 hours. Cell lysates are then obtained using Passive lysis 5X buffer (Promega; E1941). The firefly and renilla luciferase activity was determined using DualGlo luciferase assay kit (Promega; E2920) according to the manufacturer protocol. The renilla luciferase activity measurement was utilised as control for transfection efficiency and therefore the ERE_Luciferase activity was normalised to the reading obtained for the renilla luciferase activity.

### SRB assay

Briefly, the sulphorhodamine B (SRB) assay was used to monitor the effects of silencing either SLC9A3R1 or YY1, using siRNAs, on cell proliferation monolayer cultures. Cells were seeded in flat-bottomed 96-well plates (Costar; CLS3585) at a density of 2 × 10^3^. Cells were allowed to attach overnight after which, the first plate (Day 0) is assayed after the cells have become adherent. Prospective plates are assayed sequentially after 3 days, 5 days and 7 days. The cells are fixed by adding 200uL of cold 40% (weight/volume) of trichloroacetic acid (TCA) to each well for at least 60 minutes. The plates were washed five times with distilled water and then 100 uL/well of SRB (0.4% wt/vol SRB in 1% wt/vol acetic acid) reagent is added to each well and the plates are allowed to incubate for 30 minutes. The plates were then washed five times in 1% (wt/vol) acetic acid and allowed to dry overnight. SRB solubilisation was performed by adding 100 uL/well of 10 mM Tris HCl to the plates and allowed to shake for 30 minutes. Optical density was then measured using the Sunrise microplate reader (Tecan; Sunrise) at 492 nm. Cell proliferation is then calculated over the 7-day period using Day 0 as a baseline measurement.

### Enrichment scores

Overlap for ER (*in vivo*) vs enhancers and promoters were calculated by betoold intersect were the percentage overlap is calculated over the total number of regulatory regions within each bin against the concatenate ERα binding set (all ERα in all patients). For YY1, FOXA1 and ERα in MCF7, intersections were calculated using Cistrome. YY1 BEDFILEs were the consensus narrow peaks of two biological experiment, FOXA1 ChIP-seq data and ERα were obtained in house^16^. The core ERα was BEDFILE was obtained by converting the published dataset from ^43^ to HG19. The private ERα BEDFILE was obtained by iterative process to identify ERα binding unique to single patients prior to concatenation into a single file. Overlap represent the fraction of the original datasets (first dataset) overlapping with core ERα (second dataset). The TCGA luminal signature was obtained from ^38^. Each gene was extended for 20Kb upstream keeping in consideration the direction of transcription. A null gene list was generated by subtracting the TCGA luminal signature from a genome-wide gene list. Genes from the null list were extended in a similar way and enrichment was calculated by comparing the fraction of TCGA gene list with nearby binding vs. the null list. A list of estrogen target genes that do not respond to Tamoxifen was obtained from ^58^. Each gene was extended for 20Kb upstream keeping in consideration the direction of transcription. A null gene list was generated by subtracting the signature from a genome-wide gene list. Genes from the null list were extended in a similar way and enrichment was calculated by comparing the fraction of TAM resistant estrogen dependent gene list with nearby binding vs. the null list.

### RI-IHC correlation

FFPE sections for the patients used in the ChIP-seq section were retrieved from Imperial Tissue bank. Sections were stained with YY1 or SLC9A3R1 antibodies. Stained sections were divided in 20 sectors. 5 sectors with high tumor burden were scored for the number of IHC+ cells and results averaged. The number of IHC+ cells and the matched RI was analyzed using linear regression using Prism 5 (GraphPad software Inc.).

### ΔRI

See Supplementary Computational Methods.

### YY1 and SLC9A3R1 Pan cancer expression analysis

YY1 and SLC9A3R1 expression profile for matched Normal vs. Cancer samples was obtained using TIMER diff.exp option (https://cistrome.shinyapps.io/timer/). YY1 transcriptional analyses of breast cancer subtypes was performed in the Metabric Dataset (Curtis Breast) using probe ILMN_1770892 or TCGA dataset using Oncomine (https://www.oncomine.org/resource/login.html).]

### SLC9A3R1 Meta-analyses

SLC9A3R1 expression profile in drug resistant cell lines was performed by analysis of RNA-seq data from ^16^. SLC9A3R1 expression profile in MCF7 cells transfected with siRNA against ERα was performed by analysis of microarray data from GSE27473. SLC9A3R1 expression profile in additional LTED models was performed by analysis of microarray data from E-GEOD-19639. All statistical analyses were performed using Prism 5 (GraphPad software Inc.). Kaplan-Meier analysis using SLC9A3R1 expression were performed by re-analysis of 23 independent microarray datasets (KMPLOT), TCGA RNA-seq data or the combined Metabric Dataset. SLC9A3R1 transcriptional profile in breast cancer cell lines was obtained from the HPA RNA-seq dataset (http://www.proteinatlas.org/about/download). SLC9A3R1 transcriptional profile from tissues was obtained from the HPA, GTEx and FANTOM5 RNA-seq datasets (http://www.proteinatlas.org/about/download).

## Author Contribution

L.M. conceived the study. D.P., E.E performed the experiments. L.M., G.C., B.G., A.S., L.S., and P.S., developed and performed bioinformatic analyses. K.G., organised tissue collection. D.H., G.S., P.B., C.P., R.C.C., recruited patients and supplied tissues. S.S., performed pathology assessment of ChIP-seq processed samples. G.P., provided matched material. A.V. and G.P., performed IHC staining and scoring. All authors read and approved the manuscript.

## Acknowledgments

We want to acknowledge and thanks all patients and their families for the support and for donating the research samples. We thank Breast Cancer Now Tissue Bank (project TR0121), Imperial Tissue Bank and the LEGACY study for contributing tissues. The authors gratefully acknowledge infrastructure support from the Cancer Research UK Imperial Centre, the Imperial Experimental Cancer Medicine Centre and the National Institute for Health Research Imperial Biomedical Research Centre. L.M was supported by a CRUK fellowship (P64250) and Imperial Junior Fellowship (G53019). D.P was supported by a Wellcome Trust PhD studentship (103034/Z/13/Z). G.C was supported by a Marie Skłodowska Curie Training Grant (642691, EpiPredict). We acknowledge Lorna Watson, Iros Barozzi, Ylenia Perone and Jason Carrol for their constructive comments on the manuscript.

## Data Accession

H3K27ac data for all patients samples have been deposited at the ENA (http://www.ebi.ac.uk/ena) under project number XXXXXX.

## Supplementary Figures

Supplementary Figure 1. Hematoxylin-Eosin staining to evaluate tumor cellularity was carried out for each sample profiled using ChIP-seq. Only tumors with cellularity above 70% were analyzed.

Supplementary Figure 2. Summary statistics for ChIP-seq analyses. A) The number of individual peaks called using MACS 2.0 are shown for each patient. A q value of 0.01 was used in the peak calling analysis B) Saturation plots. Patients were permutated and the total number of region called after permutation is shown on the Y axis. 80% of total promoters were covered by permutating 4 patients, while similar saturation for enhancers was reached after permutating all 47 samples C) Distribution plots show the frequencies in function of Sharing Index for each regulatory region. Inset show median SI for promoters and enhancers.

Supplementary Figure 3. A) Spiking experiments show that relative enrichment for ERα binding correlates with the number of cells carrying the binding event. MCF7 stimulated or not with estradiol to induce ERα at the specified enhancers were mixed in different proportion with ERα-negative cells with similar genetic background before ERα binding was measured using ChIP-qPCR. Arrows indicate the binding site quantified using ChIP-qPCR B) Linear regression using patient-derived RNA levels and patient-derived SI. The analysis was repeated in three independent cohorts. Normal samples were analysed when available (small insets). The number in each box summarize relative slope for RN and RD elements, the colour of the box indicates the correlation coefficient R^2^.

Supplementary Figure 4. Transcription factor motif analyses of the entire promoter and enhancer imputed DHS landscapes. Motifs are ranked based on the ratio of observed/expected. Motif were filtered for a q Value of 10^−4^

Supplementary Figure 5. YY1 RNA levels from three independent RNA-seq datasets of normal tissues. Images were obtained using Protein Atlas Tools B) YY1 RNA levels from RNA-seq analysis of cancer cell lines. Images were obtained using Protein Atlas Tools.

Supplementary Figure 6. A) Chromatin landscape at the YY1 enhancer locus in breast cancer cells. The loops were obtained from Pol II ChIA-PET data (high score). The CTCF insulation perimeter was established from CTCF ChIA-PET data (ENCODE). Enhancers SI are shown at the bottom B) Meta-analysis of IHC data from Protein Atlas stained with YY1 antibody. RI for the individual tissues in indicated. Percentage of YY1 positive cells is also listed at the bottom of each image.

Supplementary Figure 7. A) YY1 expression comparing normal tissues and cancer tissues shows that YY1 median expression is significantly stronger in TCGA cancer tissues. Data were generated using TIMER (https://cistrome.shinyapps.io/timer/) (B) YY1 median expression is significantly higher in several breast cancers sub-classes compared to normal tissue. Data were obtained from Oncomine C) IGV snapshot of estradiol induced YY1 ChIP-seq and H3K27ac ChIP-seq from MCF7 cells near transcriptionally inactive genes.

Supplementary Figure 8. A-B) Silencing YY1 is sufficient to abrogate the growth of two independent cell line models of ERα breast cancer. Proliferation assays were conducted in biological triplicate. Error bars indicate 95% confidence intervals. Asterisks represent significance at P<0.05, 0.01, 0.001 and 0.0001 after 2-way ANOVA with Tukey’s post-test C) YY1 depletion leads to reduced transcription of common ERα target genes. Each experiment was performed in biological triplicates. Column and bars represent the average and SEM of all experiment. Asterisks represent significance at P<0.05, 0.01, 0.001 and 0.0001 after 1-way ANOVA with Dunnet’s post-test.

Supplementary Figure 9. A) SLC9A3R1 RNA expression in BC cell lines sensitive or resistant to endocrine therapies. RNA-seq data were obtained in house^16^ B) Meta-analysis of SLC9A3R1 expression in response to ERα depletion C) SLC9A3R1 RNA in additional oestrogen independent BC cell line models. Fold changes are calculated as ratio compared to parental endocrine sensitive BC cells. Oestrogen-dependent expression of progesterone receptor mRNA is shown as comparison. Original microarray codes are shown. Microarray were downloaded from GEO and re-analysed D) SLC9A3R1 transcriptional response to several stimuli is shown. Data were analysed using NURSA Transcriptomime tool (https://www.nursa.org/nursa/transcriptomine/index.jsf%3Bjsessionid%3DJ3hRuy3XjX6HeNr2aekh-rrTsXS-uVUXErsip0wY.nursa3) E) Kaplan Meier survival plots were calculated using three independent large datasets using SLC9A3R1 expression in the primary cancer as a classifier.

Supplementary Figure10. A) Cell lines from the Protein Atlas initiative were ranked based on SLC9A3R1 expression as profiled by RNA-seq B) SLC9A3R1 expression comparing normal tissues and cancer tissues shows that SLC9A3R1 median expression is significantly stronger in ERα-positive Breast cancer patients from TCGA. Data were generated using TIMER (https://cistrome.shinyapps.io/timer/) C) Chromatin landscape at the SLC9A3R1 enhancer locus in breast cancer cells. Looping analysis was conducted using Pol II ChIA-PET data. CTCF insulation perimeter was established from CTCF ChIA-PET data (ENCODE). H3K27ac, YY1, ERα, FOXA1 and DHS-seq data were developed in house and deposited online. ERα binding identified as Core are highlighted in the red box D) Expression analysis comparing median RNA expression values for several genes localized near the active SLC9A3R1 putative enhancer. SLC9A3R1 expression is significantly increased in BC tissues compared to normal while the expression of other genes is not affected

Supplementary Figure11 A) YY1 silencing in MCF7 is sufficient to decrease SLC9A3R1 expression. Each experiment was performed in biological triplicates. Column and bars represent the average and SEM of all experiment. Asterisks represent significance at P<0.05, 0.01, 0.001 and 0.0001 after 1-way ANOVA with Dunnet’s post-test B) Silencing SLC9A3R1 is sufficient to abrogate the growth of an endocrine therapy resistant cell line but not the parental, treatment naïve breast cancer cell line. Proliferation assays were conducted in biological triplicate. Error bars indicate 95% confidence intervals. Asterisks represent significance at P<0.05, 0.01, 0.001 and 0.0001 after 2-way ANOVA with Tukey’s post-test C-D) YY1 and SLC9A3R1 enhancer RIs are shown for all available ENCODE cell lines. MCF7 SLC9A3R1 RI is significantly different from the median value calculated on the entire population (without MCF7, green circle). RI and relative RNA levels (RPKM) are shown when available for the same cell type. Asterisks represent significance at P<0.05, 0.01, 0.001 and 0.0001 after one sample T-Test.

Supplementary Figure 12. A) SLC9A3R1 RNA levels from three independent RNA-seq datasets of normal tissues. Images were obtained using Protein Atlas Tools B) SLC9A3R1 enhancer activity was classified based on the relative RI index in each tissue. IHC meta-analysis from the Protein Atlas initiative supports the predicted heterogeneity based on enhancer activity.

Supplementary Figure 13. A) Protein Atlas sections stained with the indicated antibody were scored and ICH+ positive cells were plotted on the radial axis. Data for each patient are plotted in each corner. Two sections were examined for most patients. Green lines indicate clonal staining.

